# A novel paradigm for operant social learning in rats

**DOI:** 10.1101/2023.03.24.534145

**Authors:** Ida V. Rautio, Ella Holt Holmberg, Devika Kurup, Benjamin A. Dunn, Jonathan R. Whitlock

## Abstract

The ability to learn by observing the behavior of others is both energy efficient and brings high survival value, making it an important learning tool for many species in the animal kingdom. As such, several forms of observational learning have been documented in a myriad of species. In the laboratory, rodents have proven useful models for studying different forms of observational learning, however, the most robust learning paradigms typically rely on aversive stimuli, like foot shocks, to drive the social acquisition of fear. Non-fear-based tasks have also been developed, but these rarely succeed in having observer animals perform a new behavior *de novo*. Consequently, much less is known regarding the cellular mechanisms supporting non-fear-based types of learning, such as visuomotor skill acquisition. To address this we developed a reward-based social learning paradigm in adult rats, in which observer animals learn to tap lit spheres in a specific sequence by watching skilled demonstrators, with successful trials leading to rewarding intracranial stimulation in both observers and performers. Following three days of observation and a 24-hour delay, observer animals outperformed control animals on several metrics of task performance and efficiency, with a subset of observers demonstrating correct performance immediately when tested. This paradigm thus introduces a novel tool to investigate the neural circuits supporting observational learning and memory for visuomotor behavior, a phenomenon about which little is understood, particularly in rodents.

## Introduction

Throughout life, animals continually learn new associations and skills through direct experience. However, learning through first-hand interaction demands energy, involves trial, error, and uncertainty, and it can in some cases be dangerous or life-threatening. Social learning, on the other hand, permits individuals to gain knowledge by interacting with or observing others, often conspecifics, as to which actions to perform, which foods to eat or predators to avoid without risking consequences for themselves. The high survival value of observational learning is reflected by its phylogenetic prevalence, with various forms having been reported in taxa including mammals, birds, reptiles, fish, cephalopods and insects (Russo 1971; Galef 1976; Galef 2005; Zentall 2012; Zentall 2004; Fiorito and Scotto 1992; Laland and Williams 1997; Avargues-Weber and Chittka 2014; Loukola et al. 2017). The diversity and complexity of social learning imply that many kinds of neural architectures and systems are involved, and, while many of the relevant cellular mechanisms remain unknown, inroads have been made for some types of learning in certain species.

Some of the best progress to date has been made in rodents, which exhibit a variety of forms of social learning in the laboratory, including acquiring spatial search strategies (Leggio et al. 2003; Yamada and Sakurai 2018), learning novel behaviors to attain food (Russo 1971; Zentall and Levine 1972; Huang et al. 1983; Carlier and Jamon 2006; Jurado-Parras et al. 2012), or forming fearful associations by observing aversive conditioning in conspecifics (Bruchey et al. 2010; Jeon et al. 2010; Twining et al. 2017; Allsop et al. 2018). These and other paradigms often use food restriction, fear or other negative experiences as motivators since they provide strong survival incentives to rapidly acquire conditioned-unconditioned stimulus (CS-US) relationships. Though effective, these approaches also bring limitations. With food deprivation, for example, demonstrators eventually become sated while performing a task, which restricts the number of trials they perform before losing motivation. Fear-based paradigms, though highly effective in driving vicarious learning, rely chiefly on circuits within the amygdala, limbic system and other structures related to affective empathy (Jeon et al. 2010; Kim et al. 2012; Meyza et al. 2017; Allsop et al. 2018; Smith et al. 2021), so are ill-suited for investigating other types of learning that depend on perceptual-motor translation, like skill acquisition through observation. Thus, additional approaches are warranted to understand how different forms of observational learning occur in the brain in differing contexts and states.

Here, we present a novel sensory-motor observational learning paradigm that relied neither on food deprivation nor punishment, but instead used rewarding stimulation to the medial forebrain bundle (MFB) of both demonstrator and observer animals as reinforcement. The task followed the structure of a classic Pavlovian conditioning paradigm, in which observers viewed demonstrators tap two internally lit spheres in a particular sequence and, whenever the demonstrator performed a trial correctly (the CS), both animals received MFB stimulation (the US). Observers viewed performers in this manner for one session per day over three consecutive days, and learning was assessed the following day. The majority of observers acquired the task and significantly outperformed control animals who viewed a similar version of the task with naïve demonstrators. The absence of learning in control animals indicated that observers integrated the actions of skilled demonstrators with contiguous reward to adaptively modify their own behavior. This demonstrates that rats can learn to perform novel behavioral sequences by visual observation, and we discuss ways in which the paradigm can be further refined for future investigations into the neural substrates supporting observational visuomotor learning.

## Materials and methods

### Animal subjects

All experiments were performed in accordance with the Norwegian Animal Welfare Act, the European Convention for the Protection of Vertebrate Animals used for Experimental and Other Scientific Purposes, and the local veterinary authority at the Norwegian University of Science and Technology. All experiments were approved by the Norwegian Food Safety Authority (Mattilsynet; protocol ID # 25094). The study contained no randomization to experimental treatments and no blinding. Sample size (number of animals) was set *a priori* to at least 7 per condition to perform unbiased statistical analyses based on Mead’s resource equation. A total of 16 male Long-Evans rats (age: >12 weeks, weight: 393-525g at the start of experimental sessions) were used in this study.

Animals were handled daily from the age of 7 weeks until the start of the experiments. All rats were housed in groups in enriched cages prior to surgery, separated prior to surgery, and housed individually in plexiglass cages (45 × 44 × 30 cm) after surgery to avoid damage to implants. Animals had *ad libitum* access to food and water throughout the entire study and were housed in a temperature and humidity-controlled environment on a reversed 12h light/12h dark cycle. Training and experimental sessions all took place during the dark cycle.

Animals from the same litter were paired for behavioral experiments whenever possible, though well-trained performers were re-used for multiple observers in some cases. The roles of “observer” and “performer” were designated randomly if the animals were siblings. In cases where animals were not from the same litter, the older animal was designated “performer”.

### Surgery

Animals were anesthetized initially with 5% isoflurane vapor mixed with oxygen and maintained at 1-3% throughout the surgical procedure, typically lasting 1-2 hours. While anesthetized, they were placed in a stereotaxic frame and injected subcutaneously with Metacam (2.5mg/kg) and Temgesic (0.05mg/kg) as general analgesics. A local anesthetic (Marcain 0.5%) was injected under the scalp before the first surgical incision was made, followed by clearing the skull of skin and fascia. Using a high-speed dental drill, multiple holes were drilled in the skull and jeweler’s screws were inserted to later anchor dental acrylic (Kulzer GmbH, Germany) at the end of the surgery. A craniotomy was opened over the right hemisphere at AP: -2.8, ML: 1.7, and a single bipolar stimulating electrode targeting the MFB was gradually lowered in the brain to the level of the lateral hypothalamus (DV: 7.8-8.0). Stimulating electrodes were 13mm long twisted stainless-steel wires coated with polyimide (125μm diameter, 150μm with insulation; MS303/3-B/SPC, Plastics One, Canada). Prior to insertion, each electrode was suspended in 70% ethanol for 45-60 min, dried and rinsed with saline. Once the electrode was in place, the craniotomy was sealed with silicone elastomer (Kwik-Sil, World Precision Instruments Inc., USA), and a thin layer of Super-Bond (Sun Medical Co., Ltd., Japan) was applied to the skull to increase bonding strength between the skull surface and dental acrylic. Once the acrylic was applied and cured, sharp edges were removed with the dental drill at the end of surgery. Following surgery, animals were placed in a heated (32ºC) chamber to recover. Once awake and moving, they were returned to their home cage. If animals showed signs of pain or discomfort post-surgery, additional medication (Metacam (2.5mg/kg) and Temgesic (0.05mg/kg)) was given, and antibiotics (Baytril, 25mg/kg) were administered if wounds showed signs of infection. The testing of stimulating electrodes prior to behavioral training was postponed while animals underwent medical treatment.

### Behavioral arena and MFB stimulation set-up

Behavioral experiments were performed in a 93.5 cm x 42 cm x 50 cm clear plexiglass box divided in the middle by a perforated transparent barrier, creating two compartments (Fig. 1). The holes in the barrier allowed for visual, auditory and olfactory cues but were not large enough to allow physical contact between the animals. Two hollow spheres (ping pong balls; STIGA sports AB, Sweden) containing remotely triggered LEDs were mounted on top of hollow metal rods and positioned at the perimeter of the performer’s side of the box (Figs. 1a and 1b). The LEDs in each sphere were connected via insulated wires threaded through the metal rods to a Raspberry Pi 3 (Raspberry Pi Foundation, UK). The Raspberry Pi 3 powered and controlled when LEDs were illuminated, which cued the animals to tap the spheres.

**Fig 1.**
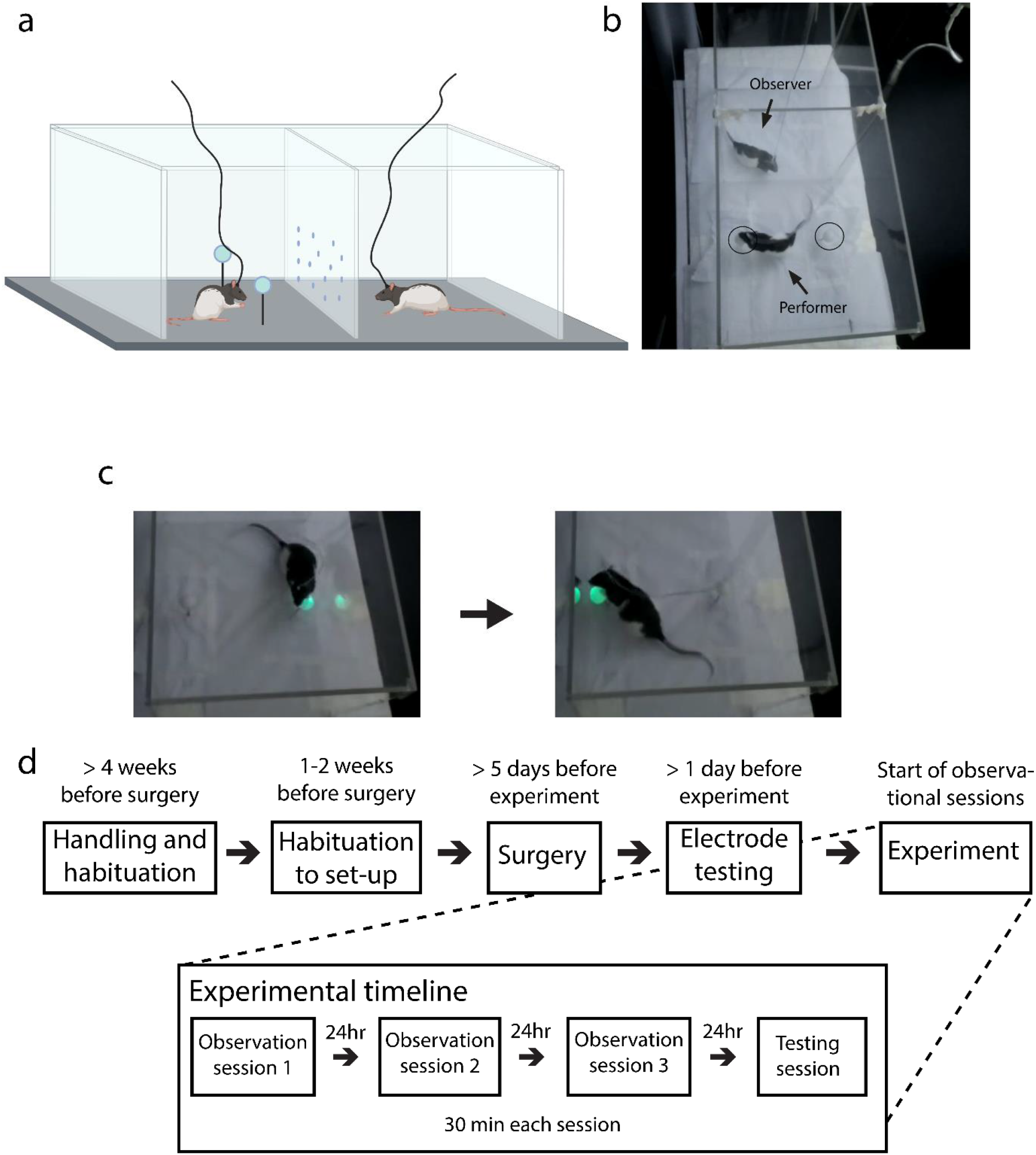
Apparatus and timeline for behavioral experiments. **a**. Schematic of the two-sided behavioral training arena, with a demonstrator animal and cue-spheres in the left chamber and an observer in the right. Schematic created with BioRender; illustration not to scale. **b**. Overhead view of an ongoing experiment, with the performer investigating an unlit cue-sphere (encircled), and the observer watching. **c**. The two-step sequence needed to attain rewarding stimulation. The picture on the left shows a performer tapping the first lit sphere, which triggers the second sphere to light up. The picture on the right shows the moment the animal taps the second sphere and receives intracranial MFB stimulation. **d**. Total timeline for the experiment, starting from before surgery to after the completion of testing

The cue and stimulation schedules in the observational learning experiments were controlled using the Raspberry Pi 3 and a standard personal Dell PC running Windows 10. Custom software written in Python 2 controlled both the LED signaling cues and delivery of electrical brain stimulation during the experiments without experimenter intervention. The Python script also allowed for manual override of the stimulation schedule, if needed, during training of performers (detailed below in Behavioral and Experimental Procedures). All training and experimental sessions lasted 30 minutes.

A Raspberry NoIR Camera V2 (Raspberry Pi Foundation, UK) was turned on each time the script was activated and recorded each session from an overhead view. The experimental room was dimly lit, and 850nm near-infrared lighting was used to illuminate the video recordings.

Intracranial stimulation of the MFB was delivered via a pulse stimulator (Master 9, Microprobes, USA) and two stimulus isolator units (SIU) (ISO-Flex, Microprobes, USA), each connected to a stimulating cable (305-305 (C)/SPC, Plastics One, Canada) connected to the head of the animal. The cord from each SIU was attached to a 2-channel commutator (Plastics One, Canada) to prevent the cords from excessive twisting while the animals moved freely in the box. Each stimulation pulse consisted of 500ms trains of square-wave pulses, each lasting 400μsec, with 200μsec on and 200μsec off per cycle, delivered at 150Hz.

## Behavioral and Experimental Procedures

### Habituation

Prior to surgery, experimental animals were handled until calm and comfortable with the experimenter and habituated to the observer-side of the apparatus, where electrodes would eventually be tested. After surgery, but before any intracranial stimulation was given, all animals were habituated to the performer-side of the box on two occasions: once without the stimulating cable attached and a second session with the cable attached. After the efficacy of the electrode was tested in the observer-side of the box (see Electrode testing, below), no further habituation or exposure to the behavioral apparatus was given until the start of the experiments. Prior to the start of any subsequent session, regardless of the stage of training or testing, each animal was allowed to habituate to their compartment until they were calm, either sitting still or grooming. After each session, the box and manipulanda were cleaned thoroughly with detergent (Zalo Ultra, Lilleborg, Norway) and wiped dry to remove trace odors or cues that might influence subsequent experimental or training sessions.

### Electrode testing

After a minimum of 5 days of post-operative recovery, stimulating electrodes were tested for efficacy and the strength of stimulation required to reinforce behavior was determined for performers and observers. Efficacy was tested by reinforcing the animal’s preference for a neutral object (*e*.*g*. a pen) in the observation chamber of the experimental apparatus. During these tests, the animal was placed in the observation-side of the box and allowed to settle (1-5 minutes), after which single stimulations were delivered when the animal oriented toward or interacted with the object. All animals started at a stimulation intensity of 20μA. If they were non-responsive then current was increased incrementally by 2μA until behavioral effects were observed, such as increased investigation or physical interaction with the object. Current was then further increased in 2μA increments until side-effects were observed (such as motor artifacts or aversive reactions), or if there had been a cumulative increase of 10μA from the identified effective stimulation intensity without any observed side-effects. If movement artifacts were elicited by the simulation, current strength was incrementally lowered by 2μA until a stimulation intensity that did not elicit artifacts was identified. The final current strength was chosen from the upper range that elicited apparent reward without side-effects (ranging from 18 μA to 60μA across animals). If the optimal range was too narrow or unclear, the animal was re-tested and the optimum was determined on a subsequent day. Electrode testing for performers took place at the start of their training (described below in *Training of performer rats*), and for observers within 24 hours before beginning the first observational session.

### Training of performer rats

The overall procedure for training demonstrator rats consisted of three phases: (i) an initial shaping phase, followed by (ii) a continuous reinforcement schedule (*i*.*e*. stimulation delivered for every sphere-tap), which then changed to (iii) a partial reinforcement phase, in which reward was delivered only when the second sphere was tapped after the first sphere had been tapped without reward. At the start of the first shaping phase, MFB stimulation was given manually whenever the animals oriented toward or approached the spheres to encourage exploration. If no behavioral changes were observed after prolonged bouts of stimulation, or if the animals showed signs of aversion, stimulation current intensity was up- or down-adjusted, respectively. If the stimulation was still aversive or had no noticeable effect, training was discontinued and the rat was excluded from the study. After the animals showed sustained interest in either of the spheres, current intensity was again fine-tuned to the lowest current strength which yielded consistent behavioral responses.

Following initial shaping, subsequent training steps were structured as follows: (1) rewarding stimulation was given when the animal started to physically interact with either sphere, (2) they were rewarded only when starting to tap each sphere alternatingly (partial reinforcement of tapping behavior), (3) rewarded when tapping the spheres as in step 2, but only when the spheres were lit (with lighting controlled manually by the experimenter), (4) withholding reward when tapping the first sphere, but delivering reward when tapping the second sphere when cued by the light, (5) fully automatic training sessions until the animal performed >75% successful trials per session. Step 5 was initiated only after the animal toggled consistently back and forth between manually lit spheres, and training was considered complete and stable once a performer exceeded 75% correct trials for three consecutive 30-minute sessions. Performer rats used for subsequent experimental sessions were given a minimum of one day of rest after reaching criterion. The automatic training sessions utilized the same script as the subsequent experimental sessions.

### Experimental task-structure

During the experiments, observer and demonstrator rats were placed in their respective sides of the box and allowed to settle. Once the animals were calm the experimenter initiated the task and an automated script turned on the LED in the first sphere with a random time-interval between 3 and 30 seconds. The first light remained on for a maximum of 30 seconds if the demonstrator did not tap the sphere. If >30 seconds elapsed, the LED turned off and the trial was scored as a “missed trial”, followed by a 3-30 second random time interval (a “time out”) before the LED turned on again and a new trial started. If the animal tapped the first sphere within the 30 seconds when the light was on, the first LED turned off and the second LED in the other sphere turned on. The second LED also had a 30 second permissive time window. If the animal tapped the second sphere within this time window, the trial was scored as “successful”, and both the performer and observer received concurrent MFB stimulation as a reward. If the demonstrator animal did not tap the second sphere within 30 seconds, the trial was scored as “failed”. After each trial, whether successful or failed, another random time-interval of 3-30 seconds elapsed before the next trial started and followed the same sequence as above. Each session lasted 30 minutes.

### Training and testing of observers

Observer animals observed either a well-trained performer or a naïve control performer (described below in Control group) for one 30-minute session per day over three consecutive days and were tested 24 hours after the final observation session. The 24-hour delay was introduced to preclude spontaneous imitation (Zentall 2006). During observational sessions, observers were able to see the performer for the entirety of each 30-minute session while allowed to move freely in their side of the box. On the day of testing, observers were placed in the performer-side of the box, and the same pre-programmed LED- and MFB stimulation-script was run as when demonstrators performed the task. No pre-training or priming with stimulations was given to the observers before testing.

### Control group

The control condition was run identically as the real experiments, but with performer animals that were completely naïve to the task. During each of the three control observation sessions, the naïve performer was allowed to move freely on the demonstrator side of the box while a custom script drove the spheres to light up and extinguish in the correct sequence, with reward stimulation delivered to both animals when the LED in the second sphere turned off. The lighting of the spheres and MFB stimulation progressed automatically, irrespective of the demonstrator’s actions, and trials were started at random and followed the same randomized 3 to 30 second time intervals as with the experimental trials. This way, reward delivery was dissociated from the behavior of the performer but maintained the sequential structure of correct trials. Critically, this condition also provided the social cue of a would-be performer but lacked the demonstration of task-specific behavior in conjunction with the reward.

### Histology

After experiments were completed, animals were deeply anesthetized with Isofluorane and injected intraperitoneally with an overdose of pentobarbital (Exagon vet., 400 mg/ml, Richter Pharma Ag, Austria), after which they were perfused intracardially with 0.9% saline and 4% paraformaldehyde. The animals were then decapitated, and skin and muscle were removed from the skull before leaving it to post-fix overnight in 4% paraformaldehyde. The following day the electrode implants were removed from the skull and the brains were extracted and stored in dimethyl sulfoxide (DMSO) at 4°C. On the day of sectioning, the brains were removed from DMSO, frozen with dry ice and sectioned with a sliding microtome at 40 μm in the coronal plane in three series. The first series was mounted immediately on glass slides, Nissl-stained and cover slipped, and the other two series were kept for long term storage in DMSO at -25°C. Electrode placement was confirmed using the Nissl-stained series (Fig. 2). Of 23 total rats implanted, two were excluded due to electrodes being off the intended target, and 5 animals had correctly placed electrodes but were excluded due to disruptive behavior (*e*.*g*. excessive jumping) on the day of testing.

**Fig 2.**
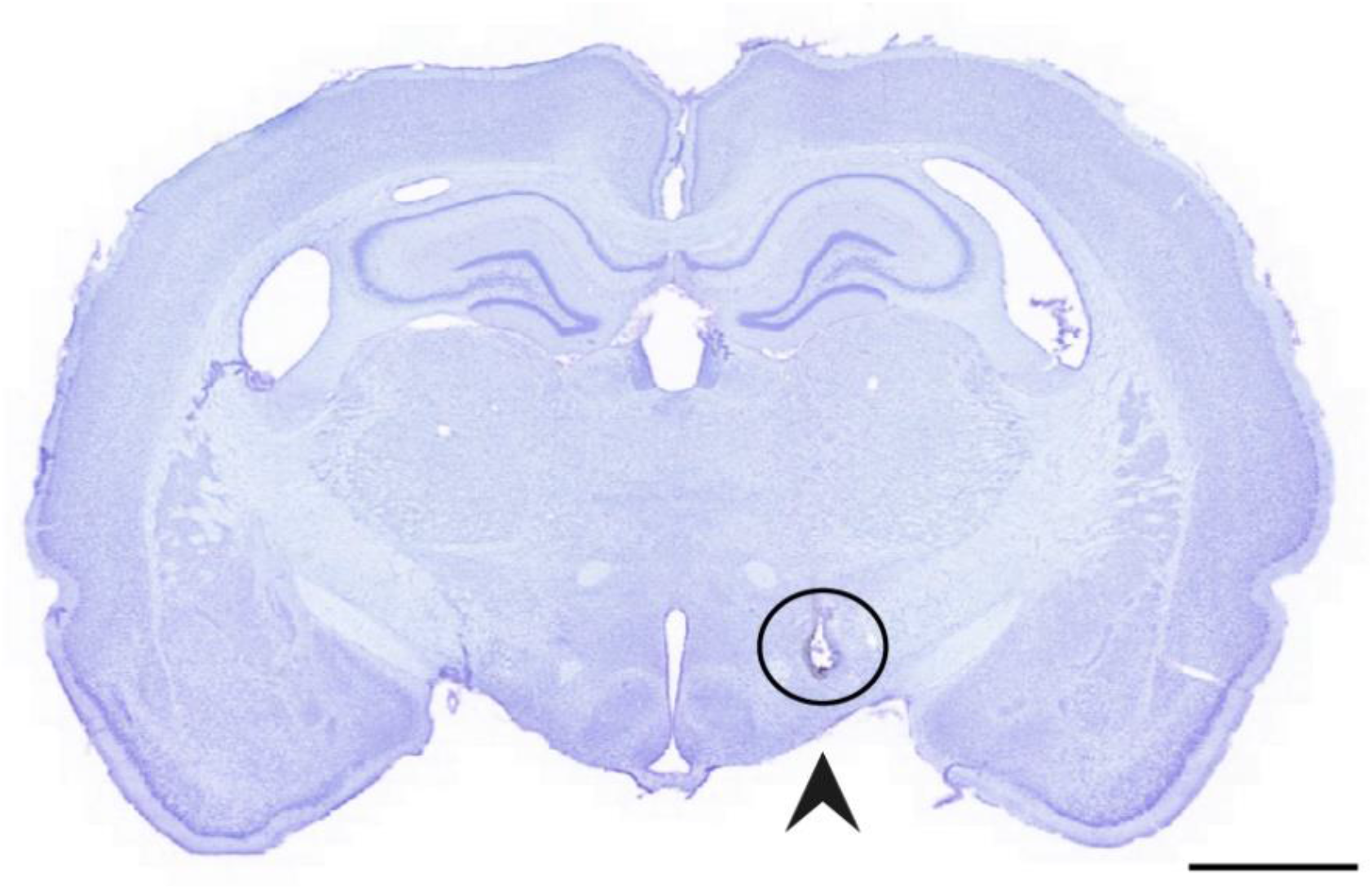
Histological verification of electrode placement. A Nissl-stained tissue section showing the termination point of a bipolar stimulating electrode targeting the medial forebrain-bundle. Scalebar = 2000 μm

## Results

Observer animals (n = 8) watched skilled demonstrators perform the sphere-tapping task in single 30-minute sessions each day for three days, with demonstrators on average performing 73.7 ± 3.0 (mean ± SEM unless otherwise indicated) trials per session, corresponding to 2.5 ± 0.1 trials per minute, and an overall success rate of 95.8%. A separate group of animals (n = 8) observed three days of the control version of the task, which consisted of naïve demonstrators, the spheres turning on and off in the correct order independently of the demonstrator’s behavior, and both observers and demonstrators receiving MFB stimulation whenever the second sphere turned off (average of 87.3 ± 0.8 automated trials per session). Importantly, the total number of MFB stimulations received during the training phase did not differ between experimental and control groups (bootstrapped independent samples t-test, *p* = 0.60). Observers and control animals were tested in the task 24 hours after the final training session, with observers performing a significantly higher proportion of correct trials (23.3 ± 8.3%) than controls (5.8 ± 3.1%; Mann-Whitney *U* = 14.5, *p* = 0.0325; Fig. 3). The rate of successful trials varied considerably across observer animals, ranging from just under 60% correct performance in the best animal to 0% in one observer who failed to learn the task (Fig. 3, left). In controls, the range of correct trials was substantially lower, with the best animal performing 23% correct trials and five with 0% (Fig. 3, right).

**Fig 3.**
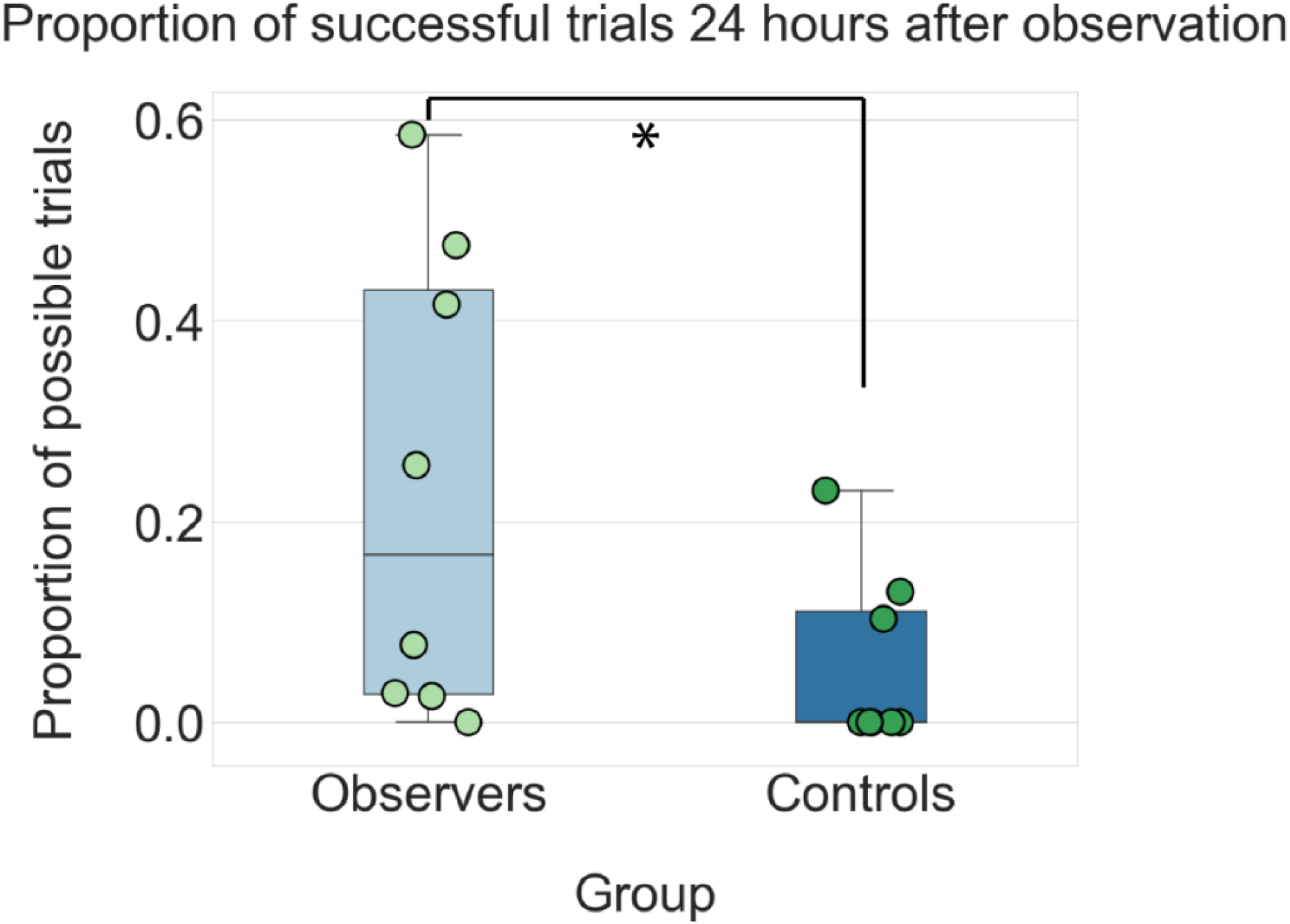
Observer animals performed a significantly higher fraction of successful trials than controls when tested 24hr after the final observation session. Whisker plots show the variability and overall success rate (out of all trials) for observers (left) and controls (right). Success rates for each animal are shown as individual dots; median performance in each group is indicated by the black line (16.7% for observers; 5.2% for controls); the 25^th^ percentile of the distribution is shown below; 75^th^ percentile is shown above; whiskers show the full distribution of data. Statistical significance between groups (*p* < .05; one-tailed Mann-Whitney U test) is noted with a star

Observer animals also exceeded controls in other aspects of task performance. These included a higher total number of correct trials performed in the test session (mean of 9.3 ± 3.3 for observers *vs*. 2.3 ± 1.2 for controls; Mann-Whitney *U* = 14.5, *p* = 0.0325; Fig. 4a), a higher number of trials performed per minute (0.30 ± 0.11 *vs*. 0.07 ± 0.04 for controls; Mann-Whitney *U* = 14.5, *p* = 0.0325; Fig. 4b), and a shorter average trial length compared to controls (45.6 ± 3.8 sec for observers vs. 54.1 ± 2.6 sec for controls; one-tailed Mann-Whitney *U* = 50, *p* = 0.0325; Fig. 4c), reflecting greater speed and efficiency in performing the task. The mean latency between tapping the first and second sphere was also shorter for observers (20.3 sec for observers *vs*. 25.5 sec for controls; Mann-Whitney *U* = 14.5, *p* = 0.0325; Fig. 4d) and, although observers had a slightly lower latency to tap the first sphere (25.5 ± 1.3 sec *vs*. 27.6 ± 0.80 sec), this did not reach statistical significance (one-tailed Mann-Whitney *U* = 19, *p* = 0.098; Fig. 4e) We note that several comparisons had the same statistical result (*i*.*e*. total successful trials, trials per minute, and mean latency between 1^st^ and 2^nd^ sphere tap) because these aspects of performance were highly correlated, resulting in similar rankings and, consequently, the same U- and p-values.

**Fig 4.**
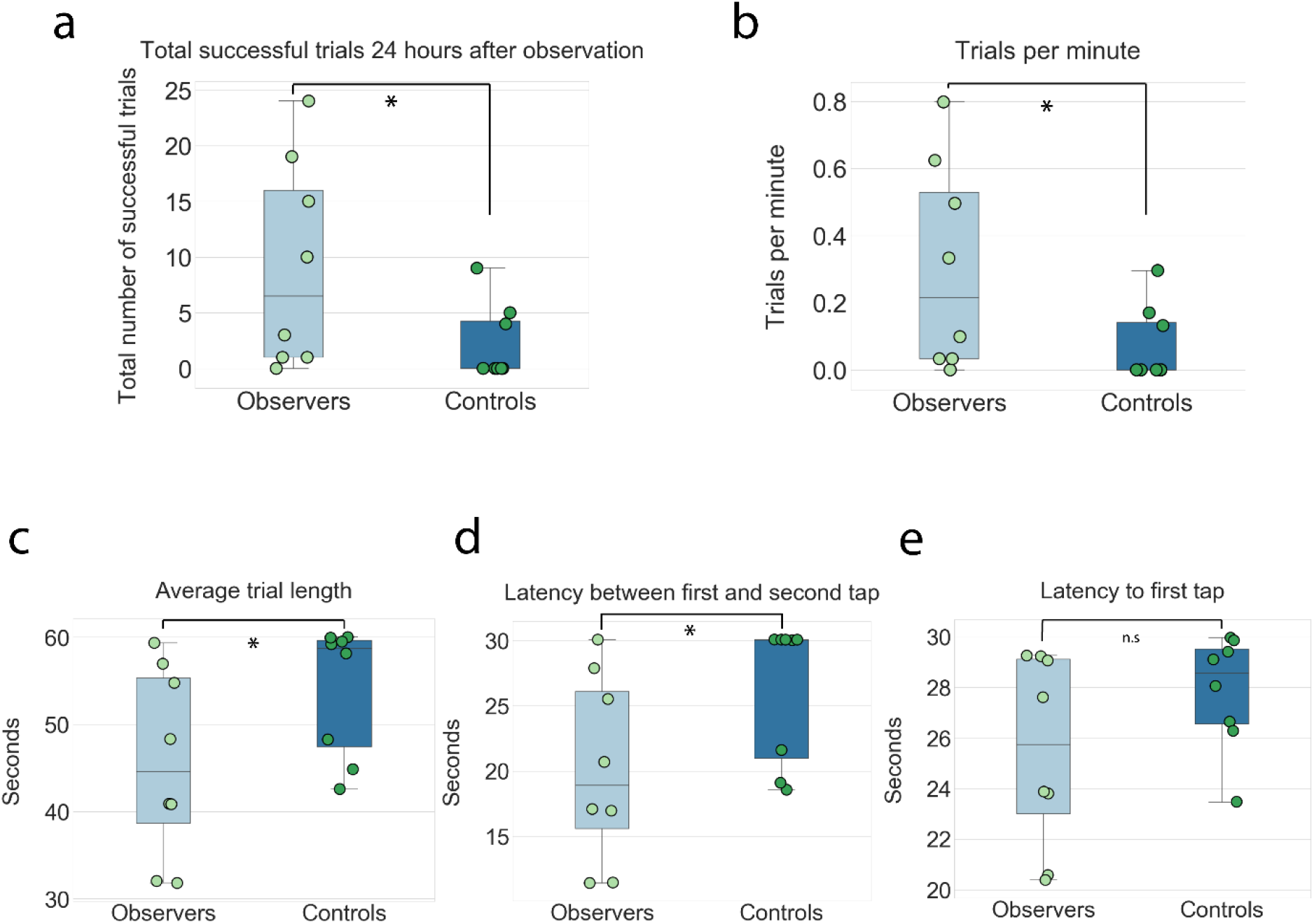
Observer animals surpassed controls on additional task performance metrics. **a**. Observers had a higher total number of successful trials than controls during testing; each dot indicates individual animal performance. **b**. Observers performed significantly more trials per minute than controls, reflecting a higher and more consistent pace of task-performance. **c**. Observers completed trials significantly faster than controls, measured as the time interval from when first sphere was lit to when the second sphere turned off; maximum possible time for each trial was 60 seconds. **d**. Observers also had a shorter average latency to tap the second sphere after the first. The maximum time for each trial was 30 seconds; with 5 of 8 control animals exceeding this time window. **e**. The average latency to tap the first sphere after it was lit was lower in observer animals, but the difference was not statistically significant. Significant differences (*p* < .05, one-tailed Mann-Whitney U test) are noted with a single star, non-significant differences are noted with n.s.

## Discussion

In this study, we demonstrate a new observational learning paradigm in which rats learn to perform a novel sequence of actions by observing the behavior of conspecifics. The task was designed in order to investigate visuomotor observational learning and its neural substrates without the use of fear, hunger or other negative experiences as motivators. The task followed the structure of a classic Pavlovian conditioning paradigm, where the demonstration of a successful trial by the performer served as the CS for observers, and rewarding intracranial stimulation served as the US. By undergoing CS-US pairing over three days, observers learned to associate task-specific actions with reward, in effect learning to perform the task by observation. The observer animals performed a higher percentage of correct trials when tested, and they outperformed controls in several temporal metrics indicating a higher efficiency in performing the task. Key design features included the use of MFB stimulation to maintain motivation and task engagement for both observers and demonstrators, as well as the use of large spherical cues that were visible from any angle. Crucially, the spheres needed to be manipulated in a specific order, such that observers had to acquire the sequence of actions to perform rather than rely on stimulus enhancement alone.

The motivation to create the task stemmed from the fact that much of our current knowledge of the neurobiological mechanisms underlying social learning derives from fear-based learning paradigms in rodents. Even though fear and pain are powerful stimuli (*e*.*g*. (Carrillo et al. 2019)) which can drive robust vicarious learning in different species (John et al. 1968; Olsson et al. 2007; Jeon et al. 2010), observational learning encompasses more than just acquisition of fearful associations, and includes other forms of learning that depend on visual, motor or spatial cognition (Heyes 1994; Galef 2005; Gariepy et al. 2014; Carcea and Froemke 2019). Naturalistic examples have been reported in a wide range of vertebrate and invertebrates, and include acquiring tool usage (Whiten et al. 1999; Biro et al. 2003; Sanz and Morgan 2007; Holzhaider et al. 2010; Loukola et al. 2017), vocal learning (Konishi 2004; Mooney 2014), learning where to shoal or forage for food (Valsecchi et al. 1989; Laland and Plotkin 1990; Laland and Williams 1997; Emery et al. 2008), or how to solve a specific task to gain access to food (Palameta and Lefebvre 1985; Zohar 1991; Prato Previde and Poli 1996; Carlier and Jamon 2006; Gruber et al. 2009). While remarkable neurobiological insights have been gained, for example, in the songbird learning system (Prather et al. 2008; Vallentin et al. 2016), progress in uncovering the neural bases of visuomotor observational learning has been impeded by a dearth of paradigms which reliably yield *de novo* skill learning in the lab. In rodents, it is more common that social learning tasks impart a general strategy or prime subjects to perform behaviors (*e*.*g*. (Leggio et al. 2003; Jurado-Parras et al. 2012)) rather than produce learners who execute a novel behavior from scratch (*e*.*g*. (Carlier and Jamon 2006)). It was therefore our goal to design a task for rodents that depended on visuomotor cognition in which at least a subset of animals performed the task correctly when first tested.

Perhaps the most crucial design aspect in the present paradigm was the use of intracranial MFB stimulation as the reward, which brought four main advantages. One was that we did not need to rely on food or water deprivation, which avoided potential deprivation-related detriments in performance, as well as the loss of motivation from demonstrators once sated. In addition, the lack of food and water restriction was better for the animals’ welfare. The second main advantage, related to the first, was that MFB stimulation was a powerful source of positive reinforcement delivered directly to the brain, bypassing the animals’ need for calories or water. It was therefore possible to sustain a high number of trials demonstrated by performers, with an average of more than 70 correct demonstrations in a 30-minute session, and in some sessions >85. This exceeded the rates of performance for demonstrators in food-motivated operant paradigms (*e*.*g*. Carlier & Jamon (2006) reported a total of 70 learning trials across 14 daily observational sessions; Jurado-Parras et al. (2012) used a lever-pressing paradigm eliciting ∼20 trials per 20-minute session). In a previous food-dependent version of the present task, we noted a steep drop in performance by demonstrators after ∼10 minutes, presumably due to the animals reaching satiety. A third advantage of MFB stimulation was the consistent salience of the reward, which may have helped sustain engagement in the task for observers as well as performers. Previous studies have sought to enhance the attention of observers by using performers of the opposite sex (Collins 1988; Carlier and Jamon 2006), or by tapping into the natural inclination of weanling animals to observe their mothers or older adults (Valsecchi et al. 1989; Prato Previde and Poli 1996). The use of weanling animals in particular proved effective, though the use of pups could present challenges for recording or manipulating the brain due to their small size. A final advantage of using MFB stimulation was that its delivery, as well as the other timed features of the task, were automated, which gave precise programmatic control over the timing of events within a trial. This reduced training variability within and between animals and allowed us to systematically adjust timing intervals (*e*.*g*. upper and lower bounds of inter-trial intervals) while developing the task.

As not all experimental animals learned the task to the same degree, with some learning poorly, there is still room for improving the paradigm. Based on earlier in-house observations, one way to better the performance of observers would be to increase the total number of observation trials by increasing from three to 5 days of training. For example, while developing the task, two pilot animals with 5 days of training gave > 91 % successful performance during testing. However, a caveat was that the animals were tested immediately after the final observational session, so their high rate of performance could have been explained as a result of spontaneous imitation (Zentall 2006). This prompted the inclusion of a 24-hour delay to remove such effects, though we note that reducing the delay to 8 or 12 hours might produce more robust performance while still testing genuine associative learning. Another way to increase the total number of observation trials would be to use session lengths longer than 30 minutes, but careful attention should be given not to fatigue the demonstrator animals. We found that well-trained performers were in constant motion for over 30 minutes, and most animals’ performance began to decline beyond this point.

By optimizing the features above to yield maximal performance over three days of training, we demonstrated the proof of principle for an observational visuomotor learning task in rats which does not require aversive stimulation or food deprivation. With potential further refinement, the task could serve as a powerful tool for studies seeking to elucidate the neural substrates supporting the acquisition and long-term (>24hr) memory of social visuomotor learning. More generally, the establishment of such a paradigm opens the door for a broader comparison of pathway-specific processes supporting different kinds of observational learning, such as those depending on visuomotor cognition *versus* fear-motivated associative learning.

## Acknowledgements

We thank M. Andresen, C. Bjørkli, K. Haugen, A.M. Amundsgård, P. J. B. Girão, and H. Waade for technical and IT assistance; S. Eggen for veterinary oversight; M. P. Witter, G.M Olsen, K. Hovde and members of the Whitlock lab for helpful discussions.

## Compliance with Ethical Standards

The authors declare that they have no conflict of interest.

## Data availability

The datasets generated during the current study are available from the corresponding author on reasonable request.

## Author contributions

Conceptualization: Jonathan Whitlock, Benjamin Dunn, Ida V. Rautio; Methodology: Ida V. Rautio, Benjamin Dunn, Jonathan Whitlock; Formal analysis and investigation: Ida V. Rautio, Ella Holt Holmberg, Devika Kurup; Project administration: Ida V. Rautio, Jonathan Whitlock; Funding acquisition: Jonathan Whitlock; Resources: Jonathan Whitlock; Supervision: Jonathan Whitlock, Benjamin Dunn, Ida V. Rautio; Writing - original draft preparation: Ida V. Rautio; Writing - review and editing: Ida V. Rautio, Jonathan Whitlock, Benjamin Dunn, Ella Holt Holmberg, Devika Kurup

## References

Allsop, S. A., R. Wichmann, F. Mills, A. Burgos-Robles, C. J. Chang, A. C. Felix-Ortiz, A. Vienne, et al. (2018) Corticoamygdala Transfer of Socially Derived Information Gates Observational Learning. Cell 173, 6: 1329–42 e18. https://doi.org/10.1016/j.cell.2018.04.004

Avargues-Weber, A., and L. Chittka (2014) Observational Conditioning in Flower Choice Copying by Bumblebees (Bombus Terrestris): Influence of Observer Distance and Demonstrator Movement. PLoS One 9, 2: e88415. https://doi.org/10.1371/journal.pone.0088415

Biro, D., N. Inoue-Nakamura, R. Tonooka, G. Yamakoshi, C. Sousa, and T. Matsuzawa (2003) Cultural Innovation and Transmission of Tool Use in Wild Chimpanzees: Evidence from Field Experiments. Anim Cogn 6, 4: 213–23. https://doi.org/10.1007/s10071-003-0183-x

Bruchey, A. K., C. E. Jones, and M. H. Monfils (2010) Fear Conditioning by-Proxy: Social Transmission of Fear During Memory Retrieval. Behav Brain Res 214, 1: 80–4. https://doi.org/10.1016/j.bbr.2010.04.047

Carcea, I., and R. C. Froemke (2019) Biological Mechanisms for Observational Learning. Curr Opin Neurobiol 54: 178–85. https://doi.org/10.1016/j.conb.2018.11.008

Carlier, P., and M. Jamon (2006) Observational Learning in C57bl/6j Mice. Behav Brain Res 174, 1: 125–31. https://doi.org/10.1016/j.bbr.2006.07.014

Carrillo, M., Y. Han, F. Migliorati, M. Liu, V. Gazzola, and C. Keysers (2019) Emotional Mirror Neurons in the Rat’s Anterior Cingulate Cortex. Curr Biol 29, 8: 1301–12 e6. https://doi.org/10.1016/j.cub.2019.03.024

Collins, R. L. (1988) Observational Learning of a Left-Right Behavioral Asymmetry in Mice (Mus Musculus). J Comp Psychol 102, 3: 222–4. https://doi.org/10.1037/0735-7036.102.3.222

Emery, Nathan, Joanna Dally, and Nicola Clayton (2008) Social Influences on Foraging by Rooks (Corvus Frugilegus). [In English]. Behaviour 145, 8: 1101–24. https://doi.org/https://doi.org/10.1163/156853908784474470

Fiorito, G., and P. Scotto (1992) Observational Learning in Octopus Vulgaris. Science 256, 5056: 545–7. https://doi.org/10.1126/science.256.5056.545

Galef, B. G., Jr.; Laland, K. N. (2005) Social Learning in Animals: Empirical Studies and Theoretical Models. BioScience 55, 6: 489–99. https://doi.org/10.1641/0006-3568(2005)055[0489:SLIAES]2.0.CO;2

Galef, B.G., Jr. (1976) Social Transmission of Acquired Behavior: A Discussion of Tradition and Social Learning in Vertebrates. In Advances in the Study of Behavior, edited by S. Rosenblatt, Hinde, R.A., Shaw, E., Beer, C., 77–100. New York: Academic Press, 1976.

Gariepy, J. F., K. K. Watson, E. Du, D. L. Xie, J. Erb, D. Amasino, and M. L. Platt (2014) Social Learning in Humans and Other Animals. Front Neurosci 8: 58. https://doi.org/10.3389/fnins.2014.00058

Gruber, T., M. N. Muller, P. Strimling, R. Wrangham, and K. Zuberbuhler (2009) Wild Chimpanzees Rely on Cultural Knowledge to Solve an Experimental Honey Acquisition Task. Curr Biol 19, 21: 1806–10. https://doi.org/10.1016/j.cub.2009.08.060

Heyes, C. M. (1994) Social Learning in Animals: Categories and Mechanisms. Biol Rev Camb Philos Soc 69, 2: 207–31

Holzhaider, J. C., G. R. Hunt, and R. D. Gray (2010) Social Learning in New Caledonian Crows. Learn Behav 38, 3: 206–19. https://doi.org/10.3758/LB.38.3.206

Huang, I. N., C. A. Koski, and J. R. DeQuardo (1983) Observational Learning of a Bar-Press by Rats. J Gen Psychol 108, 1: 103–11. https://doi.org/10.1080/00221309.1983.9711484

Jeon, D., S. Kim, M. Chetana, D. Jo, H. E. Ruley, S. Y. Lin, D. Rabah, J. P. Kinet, and H. S. Shin (2010) Observational Fear Learning Involves Affective Pain System and Cav1.2 Ca2+ Channels in Acc. Nat Neurosci 13, 4: 482–8. https://doi.org/10.1038/nn.2504

John, E. R., P. Chesler, F. Bartlett, and I. Victor (1968) Observation Learning in Cats. Science 159, 3822: 1489–91. https://doi.org/10.1126/science.159.3822.1489

Jurado-Parras, M. T., A. Gruart, and J. M. Delgado-Garcia (2012) Observational Learning in Mice Can Be Prevented by Medial Prefrontal Cortex Stimulation and Enhanced by Nucleus Accumbens Stimulation. Learn Mem 19, 3: 99–106. https://doi.org/10.1101/lm.024760.111

Kim, S., F. Matyas, S. Lee, L. Acsady, and H. S. Shin (2012) Lateralization of Observational Fear Learning at the Cortical but Not Thalamic Level in Mice. Proc Natl Acad Sci U S A 109, 38: 15497–501. https://doi.org/10.1073/pnas.1213903109

Konishi, M. (2004) The Role of Auditory Feedback in Birdsong. Ann N Y Acad Sci 1016: 463–75. https://doi.org/10.1196/annals.1298.010

Laland, K. N., and H. C. Plotkin (1990) Social-Learning and Social Transmission of Foraging Information in Norway Rats (Rattus-Norvegicus). [In English]. Animal Learning & Behavior 18, 3: 246–51. https://doi.org/Doi10.3758/Bf03205282

Laland, K. N., and K. Williams (1997) Shoaling Generates Social Learning of Foraging Information in Guppies. Anim Behav 53, 6: 1161–9. https://doi.org/10.1006/anbe.1996.0318

Leggio, M. G., A. Graziano, L. Mandolesi, M. Molinari, P. Neri, and L. Petrosini (2003) A New Paradigm to Analyze Observational Learning in Rats. Brain Res Brain Res Protoc 12, 2: 83–90

Loukola, O. J., C. Solvi, L. Coscos, and L. Chittka (2017) Bumblebees Show Cognitive Flexibility by Improving on an Observed Complex Behavior. Science 355, 6327: 833–36. https://doi.org/10.1126/science.aag2360

Meyza, K. Z., I. B. Bartal, M. H. Monfils, J. B. Panksepp, and E. Knapska (2017) The Roots of Empathy: Through the Lens of Rodent Models. Neurosci Biobehav Rev 76, Pt B: 216–34. https://doi.org/10.1016/j.neubiorev.2016.10.028

Mooney, R. (2014) Auditory-Vocal Mirroring in Songbirds. Philos Trans R Soc Lond B Biol Sci 369, 1644: 20130179. https://doi.org/10.1098/rstb.2013.0179

Olsson, A., K. I. Nearing, and E. A. Phelps (2007) Learning Fears by Observing Others: The Neural Systems of Social Fear Transmission. Soc Cogn Affect Neurosci 2, 1: 3–11. https://doi.org/10.1093/scan/nsm005

Palameta, B., and L. Lefebvre (1985) The Social Transmission of a Food-Finding Technique in Pigeons - What Is Learned. [In English]. Animal Behaviour 33, Aug: 892–96. https://doi.org/Doi10.1016/S0003-3472(85)80023-3

Prather, J. F., S. Peters, S. Nowicki, and R. Mooney (2008) Precise Auditory-Vocal Mirroring in Neurons for Learned Vocal Communication. Nature 451, 7176: 305–10. https://doi.org/10.1038/nature06492

Prato Previde, E., and M. D. Poli (1996) Social Learning in the Golden Hamster (Mesocricetus Auratus). J Comp Psychol 110, 2: 203–8. https://doi.org/10.1037/0735-7036.110.2.203

Russo, J.D. (1971) Observational Learning in Hooded Rats. Psychonomic Science 24, 1: 37–38

Sanz, C. M., and D. B. Morgan (2007) Chimpanzee Tool Technology in the Goualougo Triangle, Republic of Congo. J Hum Evol 52, 4: 420–33. https://doi.org/10.1016/j.jhevol.2006.11.001

Smith, M. L., N. Asada, and R. C. Malenka (2021) Anterior Cingulate Inputs to Nucleus Accumbens Control the Social Transfer of Pain and Analgesia. Science 371, 6525: 153–59. https://doi.org/10.1126/science.abe3040

Twining, R. C., J. E. Vantrease, S. Love, M. Padival, and J. A. Rosenkranz (2017) An Intra-Amygdala Circuit Specifically Regulates Social Fear Learning. Nat Neurosci 20, 3: 459–69. https://doi.org/10.1038/nn.4481

Vallentin, D., G. Kosche, D. Lipkind, and M. A. Long (2016) Neural Circuits. Inhibition Protects Acquired Song Segments During Vocal Learning in Zebra Finches. Science 351, 6270: 267–71. https://doi.org/10.1126/science.aad3023

Valsecchi, P., M. Mainardi, D. Mainardi, and I. Bosellini (1989) On the Role of the Demonstrator for the Solution of a Problem in the House Mouse. Ethology Ecology & Evolution 1, 2: 213–16. https://doi.org/10.1080/08927014.1989.9525524

Valsecchi, P., M. Mainardi, A. Sgoifo, and A. Taticchi (1989) Maternal Influences on Food Preferences in Weanling Mice Mus Domesticus. Behav Processes 19, 1-3: 155–66. https://doi.org/10.1016/0376-6357(89)90038-7

Whiten, A., J. Goodall, W. C. McGrew, T. Nishida, V. Reynolds, Y. Sugiyama, C. E. Tutin, R. W. Wrangham, and C. Boesch (1999) Cultures in Chimpanzees. Nature 399, 6737: 682–5. https://doi.org/10.1038/21415

Yamada, M., and Y. Sakurai (2018) An Observational Learning Task Using Barnes Maze in Rats. Cogn Neurodyn 12, 5: 519–23. https://doi.org/10.1007/s11571-018-9493-1

Zentall, T. R. (2004) Action Imitation in Birds. Learn Behav 32, 1: 15–23. https://doi.org/10.3758/bf03196003

Zentall, T. R. (2006) Imitation: Definitions, Evidence, and Mechanisms. Anim Cogn 9, 4: 335–53. https://doi.org/10.1007/s10071-006-0039-2

Zentall, T. R. (2012) Perspectives on Observational Learning in Animals. J Comp Psychol 126, 2: 114–28. https://doi.org/10.1037/a0025381

Zentall, T. R., and J. M. Levine (1972) Observational Learning and Social Facilitation in the Rat. Science 178, 4066: 1220–1. https://doi.org/10.1126/science.178.4066.1220

Zohar, O., Terkel, J. (1991) Acquisition of Pine Cone Stripping Behavior in Black Rats (Rattus Rattus). International Journal of Comparative Psychology 5, 1: 1–6

